# Python Interfaces for the Smoldyn Simulator

**DOI:** 10.1101/2020.12.15.422958

**Authors:** Dilawar Singh, Steven S. Andrews

## Abstract

**Motivation:** Smoldyn is a particle-based biochemical simulator that is frequently used for systems biology and biophysics research. Previously, users could only define models using text-based input or a C/C_++_ applicaton programming interface (API), which were convenient, but limited extensibility.

**Results:** We added a Python API to Smoldyn to improve integration with other software tools such as Jupyter notebooks, other Python code libraries, and other simulators. It includes low-level functions that closely mimic the existing C/C_++_ API and higher-level functions that are more convenient to use. These latter functions follow modern object-oriented Python conventions.

**Availability:** Smoldyn is open source and free, available at http://www.smoldyn.org, and can be installed with the Python package manager `pip`. It runs on Mac, Windows, and Linux.

**Supplementary information:** Documentation is available at http://www.smoldyn.org/SmoldynManual.pdf and https://smoldyn.readthedocs.io/en/latest/python/api.html.

## 1 Introduction

Smoldyn is a biochemical simulator that represents proteins and other molecules of interest as individual spherical particles [6]. These particles diffuse, exclude volume, undergo reactions with each other, and interact with surfaces, much as real molecules do. Smoldyn is notable for its high accuracy, fast performance, and wide range of features [4]. Users typically interact with Smoldyn through a text-based interface [2] but Smoldyn can also be run through a C/C_++_ application programming interface (API) [3] or as a module within the Virtual Cell [9] or MOOSE simulators [13].

Smoldyn’s text-based input method is relatively easy to use, but has the drawbacks of not being a complete programming language and being difficult to interface with other tools. To address these issues, we developed a Python scripting interface for Smoldyn. Python is widely used in science and engineering because it is simple, powerful, and supported by a wide range of software libraries. Also, Python code is interpreted rather than compiled, which allows for interactive use and generally reduces time between development and application. Our Python API enables Smoldyn to be run as a physics engine with other user interfaces, to be linked to complementary simulators to support multi-scale modeling, or to be run within a notebook environment such as Jupyter.

## 2 Implementation and features

Smoldyn’s Python API is assembled in three levels. At the bottom, the previously existing C/C_++_ API [3], which is written in C, provides access to most of Smoldyn’s internal data structures and functions. This API is primarily composed of functions for getting and setting Smoldyn model components, setting simulation parameters, and running simulations. The middle level, written in C_++_, uses the PyBind11 library [12] to create a Python wrapper for the C/C_++_ API, thus making all of the C/C_++_ functions and data accessible from Python. PyBind11 is a simple open-source header-only C_++_ library that was designed primarily for this task of adding Python bindings to existing C_++_ code; it takes care of reference counting, object lifetime management, and other basic utilities. The top level or “user API,” which is written in Python, exposes a set of Python classes to the user. They offer functions for building and simulating models using object-oriented programming, including standard Python features such as error handling and default parameters. They work by calling the low-level Python API, which calls the C/C_++_ API.

The classes in the user API represent key model elements. These include a “simulation” class for the entire simulated system, a “species” class for chemical species, a “reaction” class for chemical reactions, a “surface” class for biological membranes or other physical surfaces, a “compartment” class for defined volumes of space, and others. A user creates a model by creating a simulation class first and then adding components to it, such as species, surfaces, and reactions. Once it’s complete, the user tells the simulation class to run the model, typically displaying the results to a graphical output window in the process. An entire simulation is encapsulated by its own object, so users can define multiple simulations at once and even have them interact with each other. Figure 1 illustrates this design with a molecule clustering simulation, showing the Python source code, graphical output, and quantitative output.

**Fig. 1.**
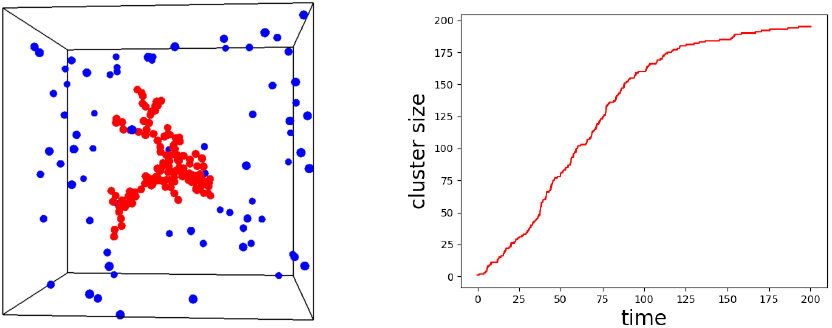
(Top) Complete Python code for a simple model of molecule clustering in which blue molecules (‘B’) diffuse freely, but then convert to immobile red molecules (‘R’) upon collision with a red molecule. (Left) A snapshot of a simulation from this model. (Right) The number of red molecules over time, from the same script.

~~~
# Smoldyn simulation of molecule clustering
import smoldyn
import matplotlib.pyplot as plt
import numpy as np
s=smoldyn.Simulation(low=[−10,−10,−10],high=[10,10,10])
B=s.addSpecies(‘B’,color=‘blue’,difc=1,display_size=0.3)
R=s.addSpecies(‘R’,color=‘red’,difc=0,display_size=0.3)
B.addToSolution(200)
R.addToSolution(1,pos=[0,0,0])
rxn=s.addReaction(name=‘r’,subs=[B,R],prds=[R,R],rate=20)
rxn.productPlacement(method=‘bounce’,param=0.6)
s.addOutputData(‘counts’)
s.addCommand(cmd=‘molcount counts’,cmd_type=‘E’)
s.setGraphics(‘opengl_good’,1)
s.run(200,dt=0.1)
data = s.getOutputData(‘counts’, 0)
dataT = np.array(data).T.tolist()
plt.plot(dataT[0],dataT[2],“r”)
plt.xlabel(“time”)
plt.ylabel(“cluster size”)
plt.show()
~~~

Figure 2 shows a study of how reduction of dimensionality reduces ligand binding times [1]. The Python API simplified this study because we could run many simulations in a row, with different adsorption coefficients, and then process and graph the data, all with a single script.

**Fig. 2.**
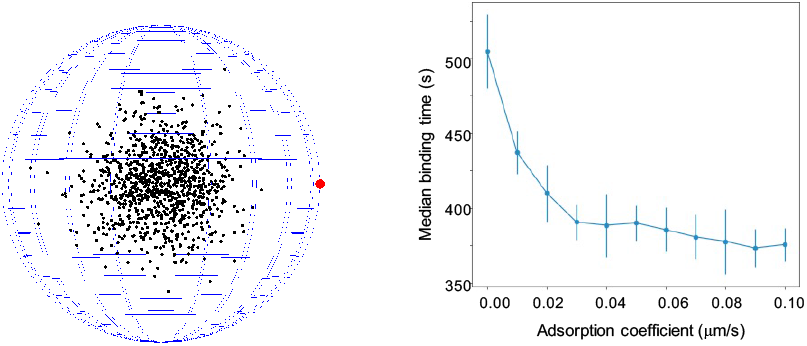
(Left) Model to investigate reduction of dimensionality effects. 1000 black molecules start at the center of a 10 *μ*m radius sphere and diffuse in the cytoplasm or on the membrane with diffusion coefficient 1 *μ*m^2^ s^−1^ until reaching the 1 *μ*m radius target, in red. (Right) Median target binding time as a function of membrane adsorption coefficient; error bars represent 1 standard deviation, for 10 runs.

As part of improving extensibility, we included callback functions in the Python user’s API. They can be used to update the Smoldyn simulation environment to states that are generated by other software, perhaps while using prior Smoldyn output to help determine that state. For example, we combined Smoldyn with the MOOSE software to simulate a simple model of pre-synaptic vesicle release. Here, MOOSE computes the membrane potential, *V_m_*, which Smoldyn then imports at every 1 ms to update the reaction rates. Internally, callbacks get registered with the Smoldyn code before a simulation starts and are then called by Smoldyn at every *n*’th cycle through the main simulation loop.

Smoldyn can compute a wide range of quantitative data during simulations, such as molecule counts in specific regions, radial distribution functions, and tracks of individual molecules. Previously, these data could only be written to text files, which could then be loaded into other software and analyzed. Now, the C/C_++_ and Python APIs also allow these data to be exported directly to other software. In both figures, for example, we transferred simulation results directly to the matplotlib graphing software.

The principal limitation of the Python API is that it does not support rule-based modeling using wildcards [5]. Additionally, Python scripts on Macintosh computers that use real-time graphical output stop executing when simulations are complete. This arises from limitations with the OpenGL graphics library versions that are available for Macs.

The Smoldyn download package includes about 15 Python scripts that demonstrate most of the Python API features. They include the three examples described above, which are called “cluster.py”, “DimensionalityEffect.py”, and “integrate_with_moose.py”.

## 3 Discussion

A particular benefit of our Python API is that it enables simple interoperability between Smoldyn and existing software libraries. We took advantage of this for the figures shown here, in both cases combining functions from Smoldyn, Numpy, and Matplotlib, where the latter two addressed data manipulation and plotting. SciPy and Pandas are other particularly useful libraries for scientific computing. With them, it would be straightforward to, for example, perform statistical inference on biochemical models using the stochastic results from well-defined Smoldyn models, adjust parameters to achieve some optimal simulation result, or simulate fluorescence microscopy images from model systems.

Several options are available for adding Python bindings to existing C/C_++_ APIs, including the Cython language [8], the SWIG automatic conversion tool [7], and Pybind11 [12]. We chose Pybind11 for several reasons. Its small size and headers-only design meant that it could be included with the main code rather than being linked, which improved software robustness and cross-platform compatibility. Also, it did not require adding an additional language to the project. Additionally, it enabled us to customize our API as desired; for example, we defined the “smoldyn.Simulation” function to accept either boundary values or an input file name, with optional arguments, and it includes a docstring with usage information.

Smoldyn is written in a combination of C, C_++_, and Python, which is partly a result of its development history, but also represents good design. The vast majority of Smoldyn’s computational effort typically occurs in core algorithms that check for molecule collisions with each other or with surfaces, and that address those collisions. We were able to make these routines fast and memory-efficient by writing them in C, which has very low computational overhead costs. The C_++_ code in the C/C_++_ API works well because of its compatibility with other C_++_ code, including PyBind11, and is a good intermediate between C and Python. Finally, the Python code in the user’s API is easy to use, fast to write and test, and an ideal interface to many other software libraries. This code is less efficient than the C code, due to Python being a high-level and interpreted language, but this has a negligible impact on total simulation times because only a tiny fraction of the total computational effort is spent here.

In some ways, Python has become the universal language of systems biology modeling because it is widely supported by a wide range of simulators, along with many high quality numerical and scientific code libraries. As a result, a single Python script can easily run multiple simulations that interact with each other. However, Python compatibility does not solve the problem of how to run a single model with different simulators because each one requires different Python code. The only viable solution is that simulators need to support systems biology standards for describing models, such as the Systems Biology Markup Language [11] for general systems biology problems, and the MUSIC language [10] for neuroscience modeling. Smoldyn does not support these standards yet but, when we add this capability, the new Python API will simplify the task.

## Acknowledgements

SSA thanks Herbert Sauro for helpful discussions and Shawn Garbett for prior work on a Python API for Smoldyn.

